# Aging impairs astrocytes in the human cerebral cortex

**DOI:** 10.1101/2022.10.31.514523

**Authors:** Alexander Popov, Nadezda Brazhe, Kseniia Morozova, Konstantin Yashin, Maxim Bychkov, Olga Nosova, Oksana Sutyagina, Alexey Brazhe, Evgenia Parshina, Li Li, Igor Medyanik, Dmitry E Korzhevskii, Zakhar Shenkarev, Ekaterina Lyukmanova, Alexei Verkhratsky, Alexey Semyanov

**Affiliations:** Department of Physiology, Jiaxing University College of Medicine, Zhejiang Pro, Jiaxing China, 314033; Shemyakin-Ovchinnikov Institute of Bioorganic Chemistry, Russian Academy of Sciences, Miklukho-Maklaya street 16/10, Moscow, 117997, Russia; Faculty of Biology, Moscow State University, Moscow, Russia; Privolzhskiy Research Medical University, Department of neurosurgery, Nizhny Novgorod, Russia; Institute of Experimental Medicine, St. Petersburg, 197376 Russia; Faculty of Biology, Medicine and Health, The University of Manchester, Manchester, M13 9PT, UK; Achucarro Center for Neuroscience, IKERBASQUE, Basque Foundation for Science, 48011 Bilbao, Spain & Department of Neurosciences, University of the Basque Country UPV/EHU and CIBERNED, Leioa, Spain; Sechenov First Moscow State Medical University, Moscow, Russia

## Abstract

How aging affects cellular components of the human brain active milieu remains largely unknown. We analyzed astrocytes and neurons in the neocortical access tissue of younger (22 - 50 years) and older (51 - 72 years) adult patients who underwent glioma resection. Aging decreased the amount of reduced mitochondrial cytochromes in astrocytes but not neurons. The total amount of protein was decreased in astrocytes and increased in neurons. Aged astrocytes showed morphological dystrophy quantified by the decreased length of branches, decreased volume fraction of leaflets, and shrinkage of the anatomical domain. Dystrophy correlated with the loss of gap junction coupling between astrocytes and increased input resistance. Aging was accompanied by the upregulation of glial fibrillary acidic protein (GFAP) and downregulation of membrane-cytoskeleton linker Ezrin associated with leaflets. No significant changes in neuronal excitability or spontaneous inhibitory postsynaptic signaling were observed. Thus, brain aging is associated with the impaired morphological presence and mitochondrial malfunction of astrocytes, but not neurons.

## Introduction

Astrocytes are crucial elements of the brain active milieu (Semyanov & Verkhratsky, 2021), being primarily responsible for homeostatic support and defense of the nervous tissue (Verkhratsky & Nedergaard, 2018). In particular, astrocytes act as a metabolic hub, storing glucose, providing metabolic substrates, and supplying neurons with glutamine obligatory for excitatory glutamatergic and inhibitory GABAergic neurotransmission (Andersen, Schousboe & Verkhratsky, 2022). Physiological brain aging is associated with a decrease in cerebral blood flow (Tarumi & Zhang, 2018) and a decline in glucose metabolism (Boemi, Furlan & Luconi, 2016). At the same time, physiological aging with preserved cognitive capacity (in contrast to neurodegeneration) is not associated with profound changes in neuron numbers and morphology (Pakkenberg & Gundersen, 1997, Verkhratsky, Mattson & Toescu, 2004). Age-dependent changes in astrocytes in the human brain are much less characterized. The numbers of astrocytes in various brain regions are not affected by old age, whereas morphometric studies are somewhat contradictory, with both increases and decreases in the size and complexity of old astrocytes being reported (Verkhratsky *et al*., 2021). Most morphological studies on astrocytes employed immunostaining with antibodies against GFAP. The GFAP immunoreactivity, however, reports neither total numbers of astrocytes nor faithfully reveals their morphology (Pekny & Pekna, 2014, Verkhratsky & Nedergaard, 2018). Studies of aged astroglia were performed almost exclusively on rodents; human astrocytes are larger and substantially more complex (Oberheim *et al*., 2009). Here we compared age-dependent changes in astrocytes and neurons in human tissue.

## Results

### Mitochondrial function is affected in aged astrocytes but not neurons

We analyzed protoplasmic astrocytes and neurons in the cortical slices prepared from the access tissue of younger adult (22 to 50 years old) and older adult (51 to 72 years old) patients of both sexes subjected to surgical resection of subcortical gliomas (Fig. 1a). Firstly, we investigated the effect of aging on the total protein content and redox state of mitochondria electron transport chain (ETC) in immunocytochemically stained astrocytes and neurons with Raman microspectroscopy (Fig. 1b). This label-free imaging technique allows monitoring vibrational modes of molecules inside cells and tissues providing information about the relative amounts, conformation, and the redox state of detectable molecules (Love *et al*., 2020, Popov *et al*., 2022). In Raman spectra, we measured peaks corresponding to lipids, proteins, and reduced cytochromes of *c* and *b*-types in the mitochondrial ETC (Fig. 1b,c). Aging decreased the relative amount of proteins in astrocytes suggesting a decline in the protein synthesis or an increase in the protein degradation rate (Fig. 1d). The amount of reduced *c* and *b*-type cytochromes in astrocytes was also lower in older adults reflecting fewer electrons in mitochondrial ETC (Fig. 1e). More pronounced decline in the reduced *c*-type cytochromes pointed to a higher rate of the electron transfer from cytochrome c to complex IV in astrocyte mitochondria in older adults (Fig. 1f). Conversely, the relative amount of proteins increased in neurons of older adults (Fig.1f). In addition, we did not observe a significant difference in reduced *c* and *b*-type cytochromes in neurons between age groups (Fig.1g,h).

**Figure 1.**
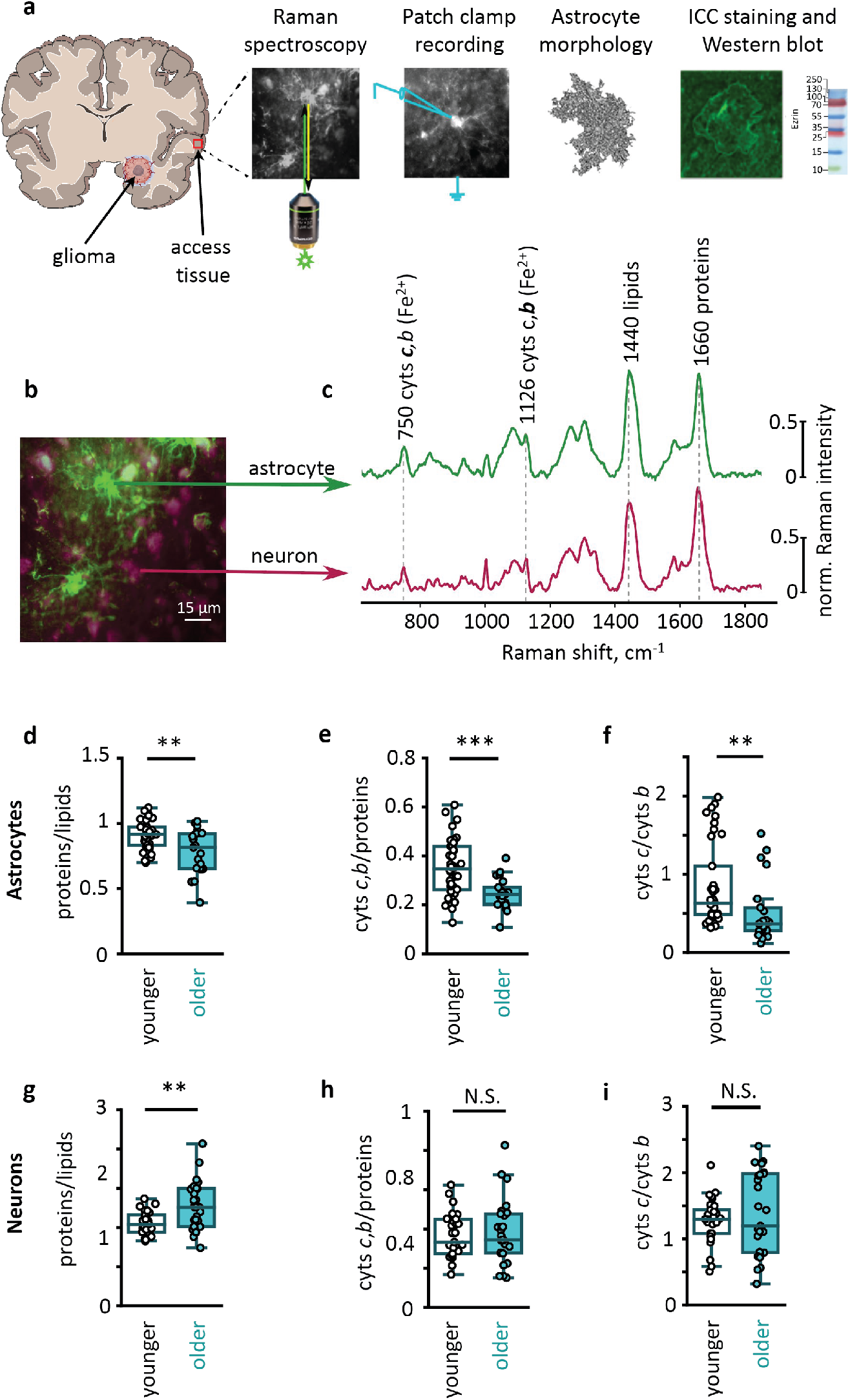
Age-dependent metabolic changes in human astrocytes and neurons. **a.** Neocortical access tissue was obtained during glioma resection. Neocortical slices were used for Raman spectroscopy, patch-clamp recordings, morphological analysis, immunocytochemical (ICC) staining, and Western blot. **b.** Double ICC-stained slice, and **c.** Raman spectra recorded from GFAP-stained astrocytes (green) and NeuN-stained neurons (burgundy). Spectral peaks marked with Raman shift value and corresponding molecule name. **d.** The ratio of proteins to lipids (p = 0.009), **e.** relative amount of reduced cytochromes of *c,b*-types (p < 0.001), and **f.** the ratio of reduced *c*-type to *b*-type cytochromes (p = 0.001) in astrocytes of two age groups (younger adults: N = 4 people, n = 38 cells; older adults: N = 3 people, n = 21 cells). **g.** The ratio of proteins to lipids (p = 0.002), **h.** relative amount of reduced cytochromes of *c*,*b*-types (p = 0.8) and **i.**the ratio of reduced *c*-type to *b*-type cytochromes (p = 0.6) in neurons of two age groups (younger adults: N = 4 people, n = 31 cells; older adults: N = 3 people, n = 27 cells) Data are shown as box-and-whisker plots where the box is Q1 and Q3 with median, whiskers are the ranges within 1.5IQR. Empty boxes/circles – younger adults, filled boxes/circles – older adults. Mann-Whitney test: N.S. – p > 0.05, ** – p < 0.01, *** – p < 0.001.

### Aging is associated with astrocytic dystrophy

Astrocytic dystrophy was reported in aged mice (Popov *et al*., 2021). We analyzed the morphology of human cortical protoplasmic astrocytes loaded with fluorescent dye Alexa Fluor 594 through a patch pipette in human cortical slices. The z-stacks of images acquired with two-photon microscopy were used for three-dimensional (3D) reconstructions of individual astrocytes (Fig. 2a). Subsequently, 3D Sholl analysis was performed on these reconstructions (Fig. 2b). We observed a decrease in astrocytic size and complexity in older human adults. Both the branch length and the maximal number of intersections decreased (Fig. 2c,d). However, neither the number of primary branches nor the ramification index was significantly affected (Fig. 2e,f).

**Figure 2.**
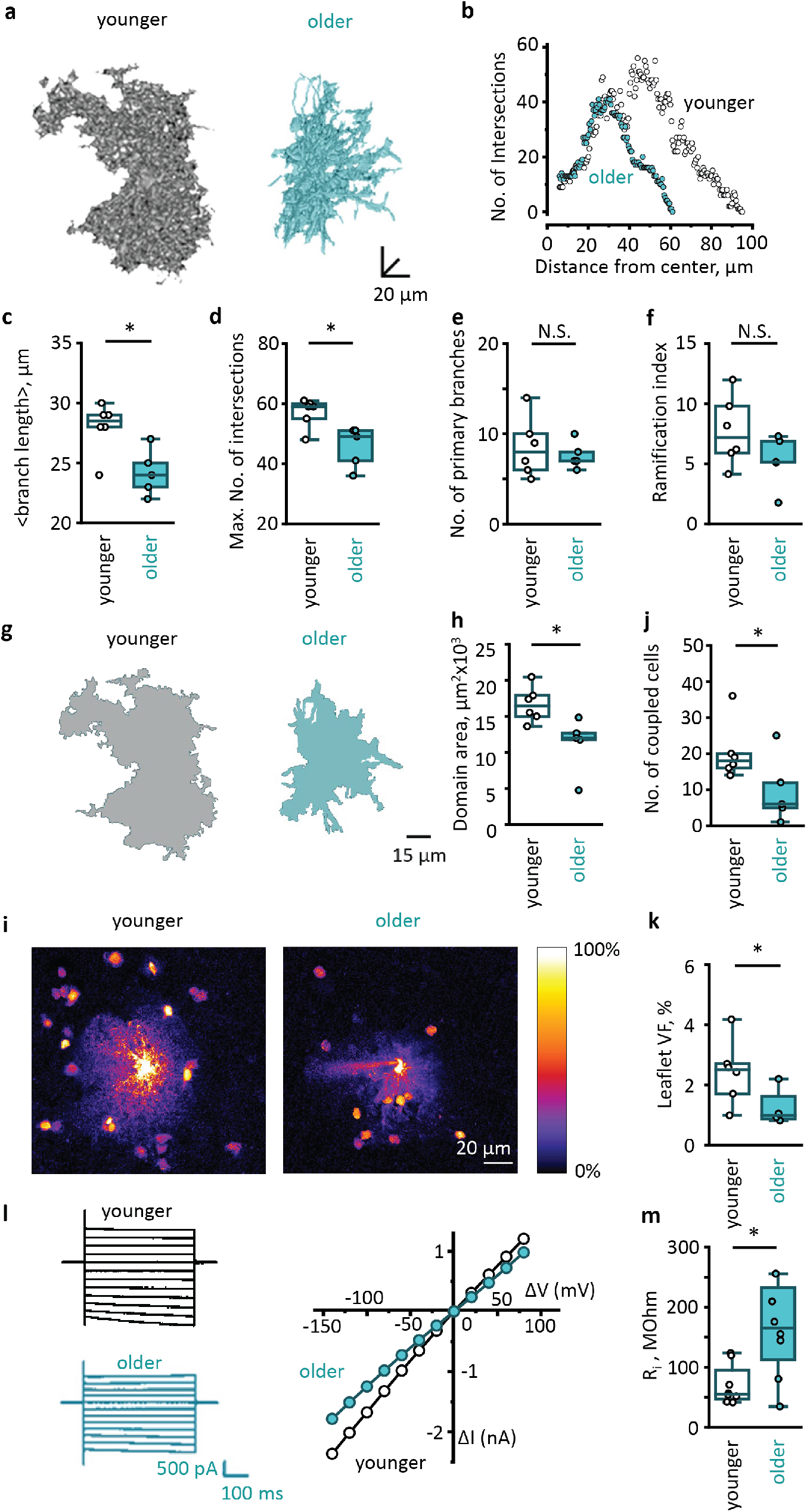
Age-dependent dystrophy of astrocytes. **a.** Representative 3D reconstructions of cortical astrocytes loaded with a fluorescent dye (Alexa Fluor 594) through patch pipette in younger (*left*) and older (*right*) adults. **b.** 3D Sholl analysis of astrocytes shown in (**a**). **c.** Mean branch lengths (p = 0.03), **d**. maximum numbers of intersections (p = 0.03), **e**. numbers of primary branches (p = 0.9) and **f**. ramification indexes (p = 0.2) in astrocytes of two age groups (younger adults: N = 6 people; older adults: N = 5 people; cell number n = N). **g.** Astrocytic domain areas obtained as z-projections of 3D reconstructions presented at (**a**) in younger (*left*) and older (*right*) adults. **h**. Astrocytic domain areas in two age groups (p = 0.01; N/n-numbers as above). **i.** Astrocytes loaded with Alexa Fluor 594 through patch pipette in younger and older adults. Note fewer neighboring astrocytes loaded through gap junctions in aged astrocyte. The fluorescence intensity was normalized to the fluorescence of soma (100%) to illustrate astrocytic spatial VF distribution. **j.** Numbers of coupled astrocytes in two age groups (p = 0.04; younger adults: N = 6; older adults: N = 5 people; cell number n = N). **k.** VFs of astrocytic leaflets in two age groups (p = 0.03; younger adults: N = 6; older adults: N = 4 people; cell number n = N). **l.** Representative currents recorded in response to voltage steps (*left*) and corresponding current-voltage relationships (*right*) in two age groups. **m.** Input resistances (Ri) of astrocytes in two age groups (p = 0.04; younger adults: N = 6 people, n = 8 cells; older adults: N = 7 people, n = 7 cells). Data are shown as box-and-whisker plots where the box is Q1 and Q3 with median, whiskers are the ranges within 1.5IQR. Empty boxes/circles – younger adults, filled boxes/circles – older adults. Mann-Whitney test. N.S. – p > 0.05, * – p < 0.05.

Then we estimated the sizes of astrocyte territorial domains by measuring the area of astrocyte image projections along the z-axis (Fig. 2g). Consistent with reduced branch length, the astrocyte domains were smaller in older adults (Fig. 2h). Smaller domains may contribute to an increase in extracellular space and uncoupling of astrocytes (Popov *et al*., 2021, Verkhratsky *et al*., 2021). Therefore, we counted the numbers of coupled astrocytes stained by fluorescent dye diffusion from the patched cell and found that the number of coupled astrocytes was smaller in older adults (Fig. 2i,j).

Protoplasmic astrocytes in the human brain contact up to 2 million synapses. These contacts are mainly formed by tiny astrocytic processes known as leaflets (Semyanov & Verkhratsky, 2021). Since astrocytic leaflets are beyond the optical resolution of the two-photon microscopy, we estimated leaflet volume fraction (VF). This method assumes fluorescence measured from astrocytic soma corresponds to 100% space occupancy. Hence, the fluorescence of unresolved astrocytic leaflets normalized to the fluorescence of soma yields the VF of the leaflets (Minge *et al*., 2021, Popov *et al*., 2022). The leaflet VF was significantly lower in older adults indicating a decrease in leaflet size, density, or both (Fig. 2k).

Shrinkage of astrocytes and loss of gap junctional coupling can affect the electrical properties of these cells (Popov *et al*., 2021, Verkhratsky *et al*., 2021). In the whole-cell voltage-clamp mode, we recorded astrocytic currents in response to depolarization steps and plotted current-voltage (IV) relationships (Fig. 2l). Shallower slop of the IV relationships in older adults indicates higher cell input resistance (Fig. 2m).

### Astrocytic dystrophy is associated with increased GFAP and decreased ezrin expression

Age-depended upregulation of GFAP is well documented (Nichols *et al*., 1993, Kohama *et al*.,1995). Based on increased GFAP immunostaining, it was concluded that astrocytes become reactive in old age (Verkerke, Hol & Middeldorp, 2021). Here, we also observed increased GFAP immunostaining in the cortex of older adults (Fig. 3 a,b). This increase in GFAP immunoreactivity in astrocytic soma and proximal branches does not contradict observed astrocyte dystrophy manifested in shrinkage of branches and leaflets that do not contain GFAP (Verkhratsky *et al*., 2021). Consistent with a previous report (Schacke *et al*., 2022), GFAP expression negatively correlated with immunostaining for membrane-to-actin cytoskeleton linker Ezrin essential for the formation and plasticity of astrocytic leaflets (Fig. 3c,d). These results were supported by Western blotting of human brain samples. In older adults, the amount of GFAP protein in the cortical tissue significantly increased (Figs. 3e,f), whereas the amount of Ezrin decreased (Figs. 3g,h). We also observed a significant increase in the protein level of glutamine synthetase (GS) in the older tissues (Figs. 3i,j).

**Figure 3.**
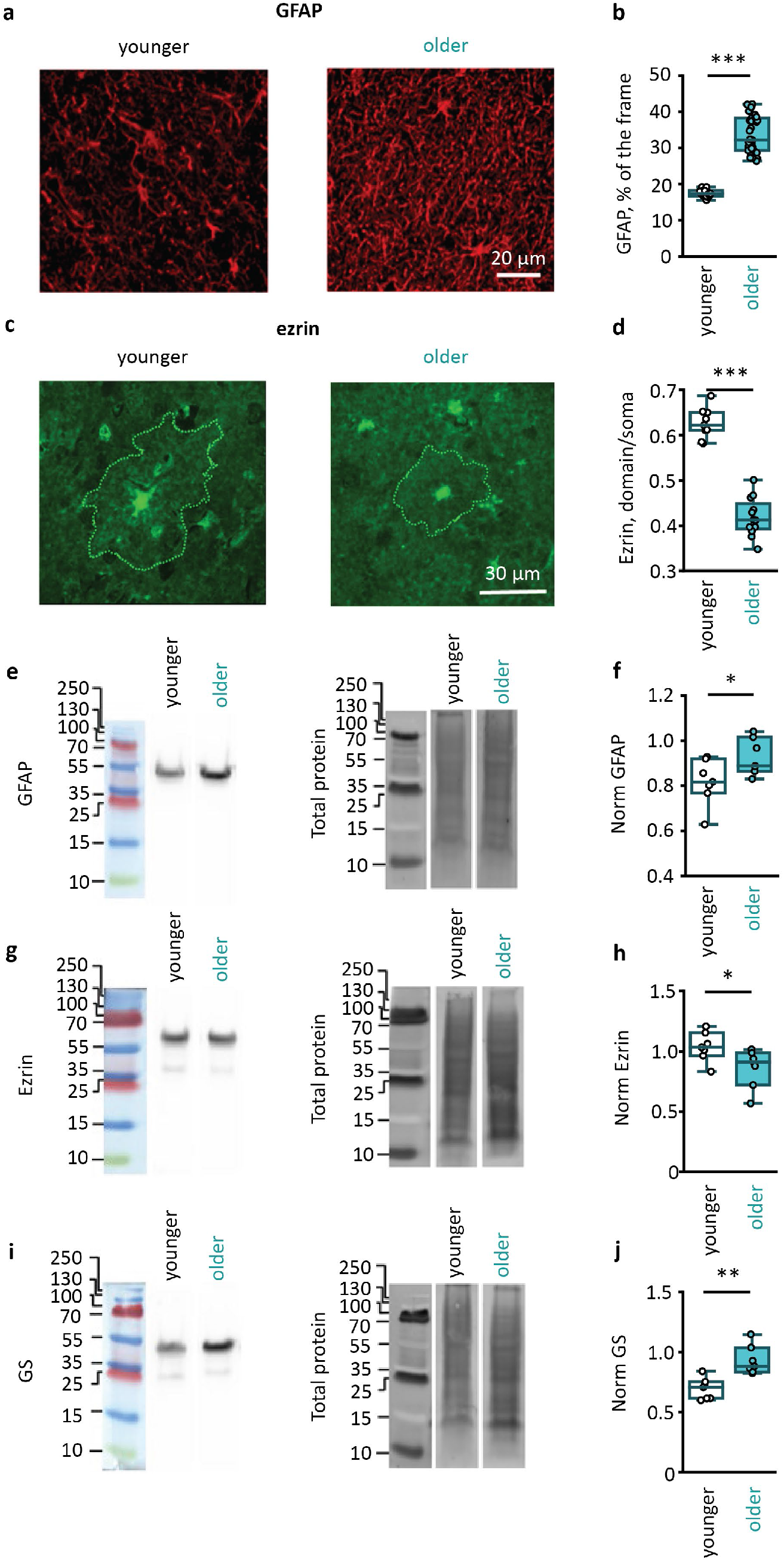
Age-dependent GFAP and GS upregulation, and Ezrin deficiency. **a.** Representative examples of immunostaining for GFAP in cortical tissue from younger (*left*)and older (*right*) adults. **b.** Percentage of the image covered by pixels stained for GFAP in two age groups (p < 0.001; younger adults: N = 3 people, n = 24 images; older adults; N = 4 people, n = 34 images). **c.** Representative examples of immunostaining for Ezrin in cortical tissue from younger (*left*) and older (*right*) adults. The astrocytic territorial domains are outlined by dotted line. **d.** Ezrin immunofluorescence averaged in the astrocyte territorial domain and normalized to the fluorescence of soma in two age groups (p < 0.001; younger adults: N = 3 people, n = 9 cells; older adults: N = 4 people, n = 12 cells).**e.** Representative Western blots of cortex homogenates stained by antibodies against GFAP (*left*) and total protein bands (*right*). **f.** GFAP amount normalized to total protein amount in two age groups (p = 0.04; younger adults: N = 7 people; older adults: N = 7 people; n = N). **g,h** same as e,f but for Ezrin (p = 0.005; N/n - numbers are the same as for f.). **i,j** same as e,f but for GS (p = 0.03; N/n - numbers are the same as for f.). Data are shown as box-and-whisker plots where the box is Q1 and Q3 with median, whiskers are the ranges within 1.5IQR. Empty boxes/circles – younger adults, filled boxes/circles – older adults. Mann-Whitney test. N.S. – p > 0.05, * – p < 0.05, ** – p < 0.01, *** – p < 0.001.

### Neuronal electrical properties and sIPSCs are not altered in the aged neocortex

We performed whole cell patch clamp recordings from layer III - IV pyramidal neurons in two age groups. Firstly, we injected hyperpolarizing voltage steps and measured the input resistance of the cells in voltage-clamp mode (Fig. 4a). No significant change in the input resistance was observed in aged neurons (Fig. 4b). Then, we injected depolarizing current steps with increasing amplitude and recorded action potentials (APs) in current clamp mode (Fig. 4c). We did not observe significant differences for any measured parameters: maximal frequency, rheobase, afterhyperpolarization (AHP), AP adaptation, AP overshoot, AP half-width between two age groups (Fig. 4d-i). These results indicate that the electrical properties of the neuronal membrane, including its excitability, do not change in the aged neocortex.

**Figure 4.**
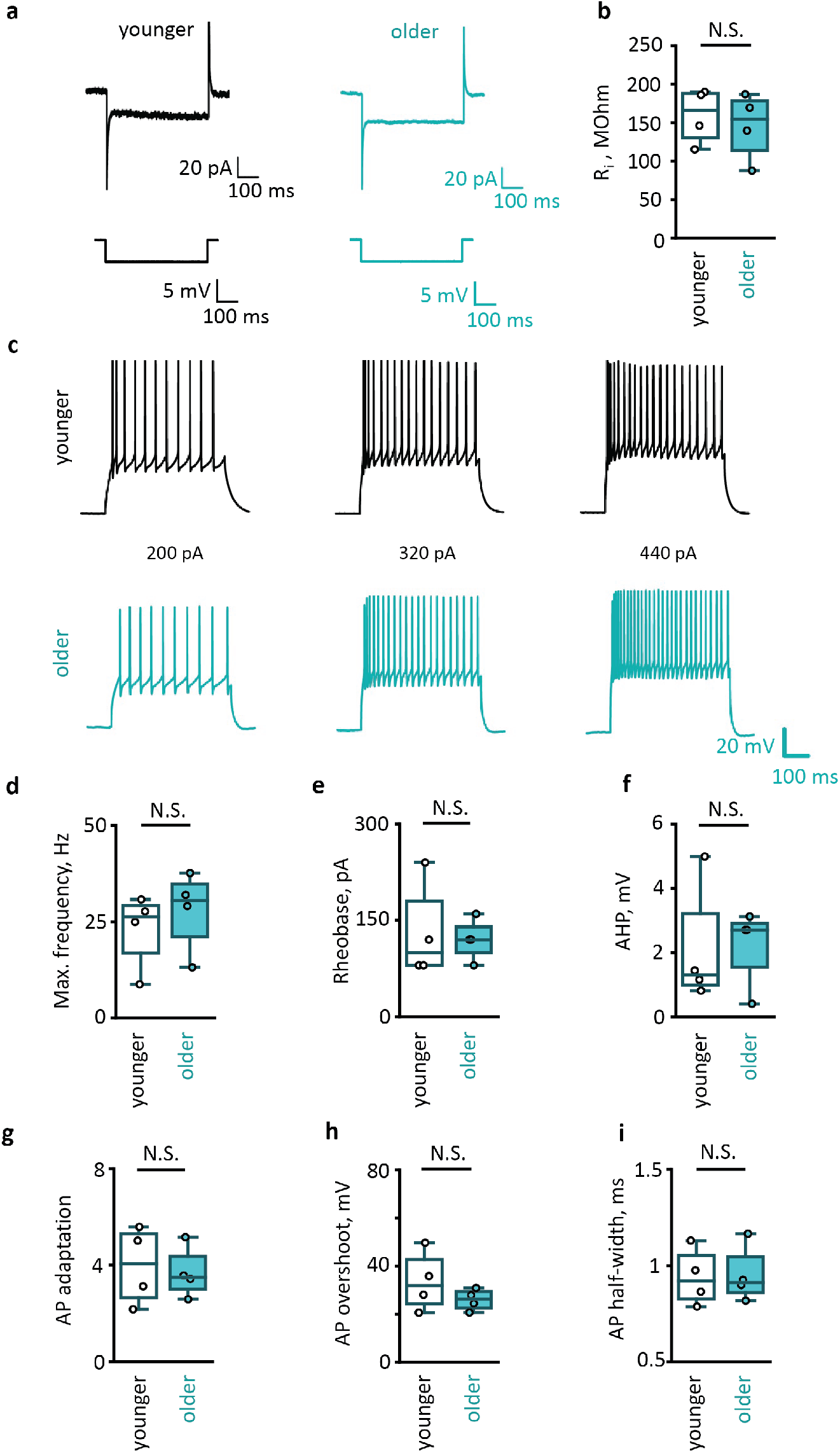
Aging does not affect membrane properties and excitability of pyramidal neurons. **a.** Representative currents recorded in cortical pyramidal neurons in response to 5 mV voltage steps in younger (*left*) and older (*right*) adults. **b.** Input resistance (Ri) of neurons in two age groups (p = 0.66; younger adults: N = 4 people; older adults: N = 4 people; cell number n = N). **c.** Action potentials (APs) recorded in current clamp mode in response to current steps 200 pA, 320 pA and 440 pA in younger (*top*) and older (*bottom*) adults. **d.** Maximal frequency of Aps (p = 0.3); **e.** rheobase (p = 0.9); **f.** afterhyperpolarization (AHP, p = 0.99); g. AP adaptation (p = 0.5); h. AP overshoot (p = 0.9); i. AP half-width in younger and older adults (N/n – numbers the same as for b.). Data are shown as box-and-whisker plots where the box is Q1 and Q3 with median, whiskers are the ranges within 1.5IQR. Empty boxes/circles – younger adults, filled boxes/circles – older adults. Mann-Whitney test. N.S. – p > 0.05.

This result does not exclude that aging affects synaptic transmission. Impairment of synaptic transmission could logically follow reduced synaptic coverage due to astrocyte dystrophy. In our experimental settings, we could monitor spontaneous inhibitory postsynaptic currents (sIPSCs) in voltage-clamp mode. However, we did not observe significant changes in sIPSC amplitude, decay, or frequency (Fig. 5).

**Figure 5.**
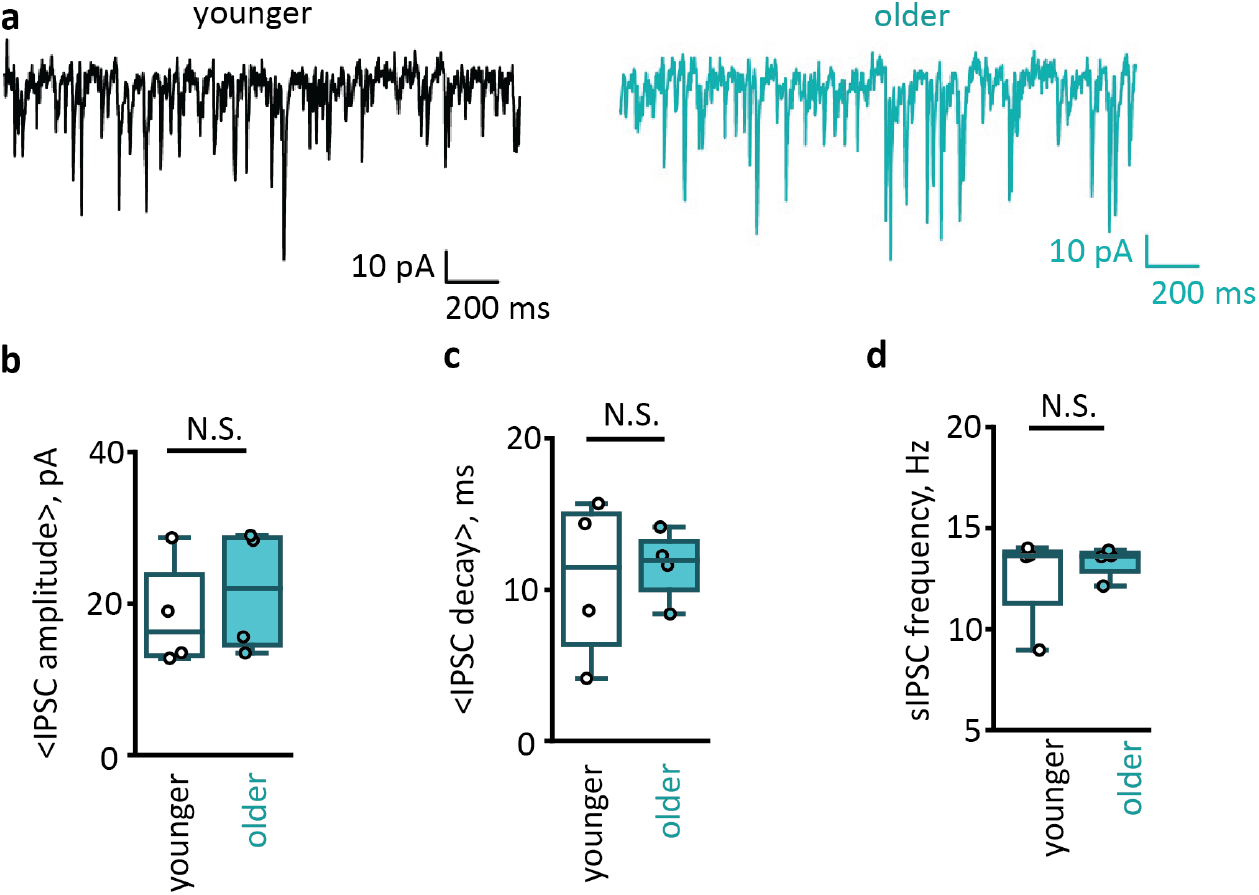
Aging does not affect sIPSCs in cortical pyramidal neurons. **a.** sIPSC recorded in voltage clamp mode in cortical pyramidal neurons. **b.** Mean amplitudes of sIPSCs in individual cells (p = 0.67), **c.** mean sIPSC decays (p = 0.89) and **d.** sIPSCs frequencies (p = 0.89) in two age groups (younger adults: N = 4 people; older adults: N = 4 people; cell number n = N). Data are shown as box-and-whisker plots where the box is Q1 and Q3 with median, whiskers are the ranges within 1.5IQR. Empty boxes/circles – younger adults, filled boxes/circles – older adults. Mann-Whitney test. N.S. – p > 0.05.

## Discussion

Single-cell Raman spectrum analysis revealed that aging is associated with metabolic changes in astrocytes but not in neurons. In particular, we found that the levels of reduced *c*- and *b*-types cytochromes were lower in aged astrocytes reflecting the decreased amount of electrons in ETC which may limit the production of mitochondrial ATP and reactive oxygen species (ROS). Mitochondrial generation of ATP is critical for astrocytic homeostatic function, as ATP fuels Na^+^/K^+^ ATPase responsible for K^+^ buffering and maintenance of Na^+^ gradients sustaining homeostatic transporters (Verkhratsky & Rose, 2020, Barros, 2022, Verkhratsky & Semyanov, 2022). ROS-signaling modulates brain metabolism and behavior (Vicente-Gutierrez *et al*., 2019). Reduced ATP and ROS supply may affect critical astrocytic support of neuronal excitability and neurotransmission.

Aging leads to morphological dystrophy of cortical protoplasmic astrocytes: the length of astrocytic branches decreases, and the territorial astrocytic domain shrinks. This observation may explain an increase in extracellular space with age (Salminen *et al*., 2016, Verkhratsky *et al*., 2021). Astrocytic morphological changes are fundamental for the regulation of extracellular space (Walch & Fiacco, 2022). An increase in extracellular space can also affect the extracellular diffusion of neurotransmitters, neuromodulators, and ions, as well as the operation of the glymphatic system (Benveniste *et al*., 2019). Aging is associated with a substantial reduction of the VF of astrocytic leaflets. Age-related decrease in VF can reflect a lower number of leaflets, smaller sizes, or both. The precise mechanism requires direct measuring of leaflet sizes and numbers. Smaller leaflet density can occur because reduced branches surface is available to carry the leaflets in older adults. Nevertheless, the lower density of leaflets indicates a reduction of astrocytic interaction with other components of the brain active milieu, including synapses (Semyanov & Verkhratsky, 2022). Synapses in the old brain therefore, have less astrocyte associations, which may limit homeostatic support, facilitate the spillover of neurotransmitters and affect synaptic plasticity (Valtcheva & Venance, 2019).

Along with astrocytic morphological dystrophy, we observed increased astrocyte input resistance. This increase most likely reflects a decrease in the membrane surface of the astrocytes, translating into a lower total membrane conductance. In addition, shrinkage of astrocytes was associated with the uncoupling of astrocytes through gap junctions. Similar uncoupling was observed in aged mice (Popov *et al*., 2021). Loss of gap junction can also increase cell input resistances, and compromise astrocyte isopotentiality within syncytium (Ma *et al*., 2016). Such age-dependent changes in electric properties can significantly affect astrocyte function. For example, potassium released during synaptic transmission depolarizes perisynaptic astrocytic processes and suppresses voltage-dependent glutamate uptake (Tyurikova *et al*., 2022). Cell input resistance defines membrane time and length constants, thus modulating the temporal and spatial properties of the glutamate uptake in the vicinity of active synapses. An increase in astrocyte input resistance prolongs and spreads astrocytic depolarization reducing glutamate uptake in old age.

Astrocytic dystrophy was associated with an increase in the expression of GFAP. Morphological analysis based on GFAP immunostaining is generally accepted as a sign of astrocytic hypertrophy and reactivity (Escartin *et al*., 2021). On the other hand, GFAP immunoreactivity is not detected in all astrocytic branches and is absent in leaflets (Semyanov & Verkhratsky, 2021). Using astrocyte staining with fluorescent dye, we revealed entire cell morphology and observed morphological dystrophy in older adults. Moreover, GFAP upregulation can trigger a deficiency of Ezrin (Schacke *et al*., 2022). Since Ezrin is essential for the formation of leaflets, downregulation of this protein may be responsible for their loss or reduction in size. Furthermore, the upregulation of GFAP can be linked to age-related astrocyte uncoupling, which is also detrimental to learning and memory (Hosli *et al*., 2022). Aging was also associated with a significant increase in astrocytic expression of GS that converts glutamate and ammonium to glutamine. Upregulation of GS may serve to reduce the toxicity of ammonia, levels of which increase in the aged brain (Jo *et al*., 2021). It may also signal some adaptive changes in the glutamate (GABA) glutamine shuttle (Andersen, Schousboe & Verkhratsky, 2022).

Notably, the mitochondrial malfunction in aged astrocytes was not accompanied by similar changes in neurons. Moreover, we observed a decrease in protein amount in astrocytes and protein accumulation in neurons. Accumulating misfolded and damaged proteins in the brain were proposed as a determinant of brain aging (Vilchez, Saez & Dillin, 2014). Nonetheless, no significant changes in basic excitability and membrane properties were observed in the neurons of older adults. No significant changes in sIPSC amplitude, decay, and frequency were observed either. Thus, impaired protein clearance may not immediately affect neuronal function. The relatively preserved neuronal function does not exclude that astrocytic atrophy is not affecting synaptic transmission and plasticity during physiological brain activity. Diminished presence of astrocytic leaflets in the brain active milieu is likely to compromise neurotransmitter uptake and potassium clearance (Popov *et al*., 2021). Extracellular accumulation of potassium could affect neuronal excitability, presynaptic release of glutamate and its uptake by astrocytes (Shih *et al*., 2013, Tyurikova *et al*., 2022). We conclude that astrocytic rather than neuronal malfunction instigates age-dependent cognitive decline in humans.

## Methods

### Human cortical slices

The specimens of access tissue were obtained during tumor resection from 42 patients of both sexes in the range of ages from 22 to 72 years. A surgical approach to the tumor was performed using a frameless navigation system with uploaded functional MRI data and intraoperative neurophysiological monitoring (as well ‘awake’ surgery) for the preservation of motor and speech eloquent brain areas and white matter tracts (Zolotova *et al*., 2022). During the resection, a fragment of cortical tissue was removed to get access to the tumor. The tissue was collected from the access area outside the area of changes on the T2-FLAIR MRI. Cortical access tissue was put in a cutting solution containing (in mM): 85 NaCl; 2.5 KCl; 26 NaHCO_3_ 1 NaH_2_PO_4_; 7 MgCl_2_; 0.5 CaCl_2_ and 50 sucrose. The solution had an osmolarity of 295 ± 5 mOsm and pH of 7.4 when saturated with 95% O_2_ and 5% CO_2_. The study was approved by the Ethical Committee of the Privolzhsky Federal Research Medical Centre of the Ministry of Health of the Russian Federation, and informed consent from patients were obtained.

Cortical access tissue was cut into 350 μm slices with a vibrating microtome HM650 V (Thermo Fisher Scientific, Waltham, USA) in the cutting solution. After preparation the slices were incubated for recovery during 1 hour at 32–34°C in a storage solution containing (in mM): 92 NaCl; 2.5 KCl; 30 NaHCO_3_ 1.2 NaH_2_PO_4_; 1 MgCl_2_; 1 CaCl_2_; 5 Na-ascorbate; 3 Na-pyruvate; 2 HEPES; 2 Thiourea and 25 D-glucose. The solution had an osmolarity of 295 ± 5 mOsm and pH of 7.4 and was saturated with carbogen (95% O_2_ and 5% CO_2_). For electrophysiology recordings and two-photon imaging, the slices were placed in an immersion chamber where they were continuously superfused (1 – 3 ml/min) with constantly carbogenized solution containing (in mM): 127 NaCl; 2.5 KCl; 1.25 NaH_2_PO_4_; 1 MgCl_2_; 2 CaCl_2_; 25 NaHCO_3_ and 25 D-glucose. The solution had an osmolarity of 295 ± 5 mOsm, pH of 7.4, and temperature of 34°C.

### Raman microspectroscopy

Cortical slices were immunocytochemically double stained for GFAP and NeuN to identify, respectively, astrocytes and neurons for subsequent Raman microspectroscopy. The staining was performed in the following steps: (1) acute slices after the recovery period were placed into 4% paraformaldehyde (PFA) solution (37°C) for 60 min and then washed twice in phosphate buffer solution (PBS, pH 7.4); (2) PFA-fixed slices were transferred into PBS with 0.3% Triton-X100 solution for 15 min and then into PBS with 0.1% Tween20 and 5% BSA solution (25°C) for 90 min; (3) the slices were incubated in the solution of primary antibodies: anti-glial fibrillar acidic protein (GFAP) rabbit polyclonal antibody (Abcam, Cambridge, UK, catalog number ab7260) and anti-neuronal nuclear protein (NeuN) chicken polyclonal antibody (Novusbio, Centeennial, USA, catalog number NBP2-104491) in PBS with 0.01% Tween 20 for 60 h at 25°C; (4) the slices were washed three times in PBS, 10 min, at 25°C; (5) the slices were incubated in the solution of secondary antibodies: CyTM2 AffiniPure donkey anti-rabbit IgG (H+L) (Jackson ImmunoResearch, Ely, UK catalog number 711-225-152) and CyTM5 AffiniPure donkey anti-chicken IgG (H+L) (Jackson ImmunoResearch, catalog number 703-175-155) for 2 h at 25°C; (5) the slices were washed twice in PBS and in trice in deionized water. Then the slices were placed on the glass slide and dried in the dark. Stained slices were stored at room temperature in the dark. The secondary antibodies were chosen according to the spectral properties of their fluorophores (excitation and emission wavelengths, CyTM2: λ_ex_ = 492 nm; λ_em_ = 510 nm and CyTM5: λ_ex_ = 650 nm; λ_em_ = 670 nm) so they did not emit fluorescence upon 532 nm laser excitation used for Raman spectroscopy.

Raman spectra were recorded with confocal Raman microspectrometer NTEGRA SPECTRA (NT-MDT, Zelenograd, Russia) attached to the inverted Olympus microscope. Astrocytes and neurons were identified by CyTM2 and CyTM5 fluorescence, respectively. Then, Raman spectra were recorded from the somas of the identified cells with the x40 NA 0.45 objective following laser excitation (532 nm, 1 mW). The spectrum accumulation time was 60 s. The laser excitation at 532 nm produced negligible CyTM2 or CyTM5 fluorescence without interfering with the Raman scattering of the stained cells. The total number of studied identified cells was: 60 astrocytes (7 patients) and 58 neurons (7 patients).

Raman spectra were analyzed with the open-source software Pyraman, available at https://github.com/abrazhe/pyraman. The baseline was subtracted in each spectrum. The baseline was defined as a cubic spline interpolation of a set of knots, numbers, and x-coordinates,which were selected manually outside any informative peaks in the spectra. Once chosen, the number and x-coordinates of the knots were fixed for all spectra in the study. Y-coordinates of the knots were defined separately for each spectrum as 5-point neighborhood averages of spectrum intensities around the user-specified x-position of the knot. The parameters for baseline subtraction were chosen after processing approximately 50 spectra from different astrocytes and neurons. After the baseline subtraction, the intensities of peaks with the following maximum positions were defined: 750, 1126, 1440, and 1660 cm^−1^. Peaks at 750 and 1126 cm-1 correspond to bond vibrations in hemes of reduced cytochromes of *c*- and *b*-types, with the main contribution from *c*-type cytochromes for the first peak and the main contribution from *b*-type cytochromes for the second peak. Peaks at 1440 and 1660 cm-1 correspond to vibrations of C-C bonds in lipids and peptide bonds in proteins, respectively (Brazhe *et al*., 2012). Relative Raman peak intensities were used to obtain the protein amount normalized to the lipid amount (I_1660_/I_1440_ ratio), the amount of reduced *c*- and *b*-type cytochromes (with the main contribution from reduced c-types cytochromes) normalized on the protein amount (I_750_/I_1660_ ratio), the ratio of reduced *c*-type cytochromes to reduced *b*-type cytochromes (I_750_/I_1126_ ratio) in astrocytes and neurons.

### Electrophysiological recordings

Astrocytes were selected at the border of cortical layers II - III. Whole-cell recordings were performed with borosilicate glass pipettes (5 – 7 MΩ) filled with an internal solution containing (in mM): 135 KCH3SO3, 10 HEPES, 10 Na2phosphocreatine, 8 NaCl, 4 Na2ATP and 0.4 NaGTP (pH was adjusted to 7.2 with KOH; osmolarity to 290 mOsm). 50 μM Alexa Fluor 594 (Thermo Fisher Scientific, USA) was added to the solution to reveal cell morphology. Passive astrocytes were identified by strongly negative resting membrane potential and linear current-voltage (IV) relationship. In the current-clamp mode, current steps were applied to corroborate the absence of membrane excitability. In voltage-clamp recordings, the astrocytes were held at −80 mV. Voltage steps (Δ20 mV; 500 ms) from −140 mV to +80 mV were applied to obtain IV relationships. All responses were amplified with a Multiclamp 700B amplifier (Molecular Devices, San Jose, USA), digitized with digital-analog converter board NI PCI-6223 (National Instruments, Austin, USA), and recorded with WinWCP v5.2.5 software (University of Strathclyde, UK). The data were analyzed with the Clampfit 10.4 software (Molecular Devices, USA).

Neurons were selected in cortical layer III. Whole-cell recordings were performed with borosilicate glass pipettes (4–5 MΩ). Cell excitability was measured with a pipette solution containing (in mM): 140 K-gluconate, 8 NaCl, 0.2 CaCl2, 10 HEPES, 2 EGTA, 0.5 NaGTP, and 2 MgATP (pH was adjusted to 7.2 with KOH and osmolarity to 290 mOsm). Membrane potential was manually set at −70 mV in current-clamp mode. We recorded cell spiking in response to 500-ms current steps of increasing amplitudes from 0 pA to 440 pA in steps of 40 pA. The maximum frequency of action potentials was calculated in response to the current step of 440 pA. The rheobase was calculated as the amplitude of the previous current step, after which action potentials appeared. The afterhyperpolarization (AHP) was calculated as the amplitude of a negative peak appearing after depolarizing step relative to the baseline before the depolarizing step. We calculated the spike adaptation rate as a ratio of the instantaneous frequency of the first and the second AP to the instantaneous frequency of the last and previous AP in response to 440 pA current step. AP overshoot was calculated as AP peak value over 0 mV. AP half-width was calculated as the duration of the action potential at the voltage halfway between the threshold and the action potential peak. Cell input resistance was calculated in voltage-clamp mode from the current in response to 5 mV depolarizing step.

GABA_A_ receptor-mediated sIPSCs were recorded in voltage-clamp mode (holding potential −70 mV) in the presence of 25 μM NBQX, 50 μM APV, and 5 μM CGP52432 to block AMPA, NMDA, and GABAB receptors, respectively. The recordings were done with intracellular solution containing (in mM): 135 CsCl, 10 HEPES, 10 phosphocreatine, 4 MgATP, 0.3 TrisGTP, 0.3 EGTA (pH 7.35 was adjusted with CsOH). sIPSCs were identified using a semi-automated detection software Mini Analysis 6.0.7 (Synaptosoft, Fort Lee, NJ, USA). The amplitude, frequency, and decay time constant of sIPSCs were measured. sIPSC amplitude and decay were averaged in individual sweeps.

### Astrocyte morphological analysis

Astrocytes were filled with 50 μM Alexa 594 through a patch pipette. Then we waited for at least 10 min to allow for sufficient dye diffusion in the astroglial syncytium. Astrocyte imaging was performed with a two-photon laser scanning microscope Zeiss LSM 7 MP (Carl Zeiss, Germany) equipped with femtosecond laser Chameleon Vision II (Coherent, UK). Alexa 594 was excited at 830 nm. Z-stacks of 100 images (512 × 512 pixels size, 0.2 μm/px) were collected with 1 μm steps. Z-axis increments were next re-sampled by spline interpolation to match the lateral resolution of 0.2 μm/px. For astrocytes morphometry, we used Image-funcut [image-funcut, https://github.com/abrazhe/image-funcut], Scikit-Image [scikit, http://scikit-image.org/], and Sci-Py [scipy, http://www.scipy.org/] libraries (van der Walt *et al*., 2014, Virtanen *et al*., 2020).

Astrocyte branches were traced with the Simple neurite tracer plugin in ImageJ (Longair, Baker & Armstrong, 2011). Then, 3D Sholl analysis was performed: the center of the soma was taken as the center for the set of concentric spheres with increasing radius (from 10 to 100 μm with a step of 1 μm). We collected the numbers of individual astrocytic branches intersected by each sphere to build the Sholl profile: a graph of the number of intersections versus the distance from the center of the soma. The number of primary branches, the maximal number of intersections, and the ramification index (the ratio of maximal intersection to primary branches) were calculated from the Sholl analysis. In addition, we calculated the mean branch length and the astrocyte domain area (the area circumscribed by a line connecting endpoints of the traced astrocyte processes in astrocyte maximal projection).

Next, we chose the frame within the Z-stack containing astrocyte soma. Special attention was paid that the fluorescence of soma was not saturated (Popov *et al*., 2020). We constructed five fluorescence cross-sections passing through the center of the soma (72° from each other). We discarded large fluctuations (> 10% and >0.5 μm) of fluorescence corresponding to astrocytic branches. Next, we estimated the volume fraction (VF) of astrocytic leaflets as relative fluorescence intensity in the unresolved processes normalized to fluorescence intensity in the soma. Characteristic VF was obtained at a distance of 30–40 μm from the soma boundary.

### Immunocytochemical analysis

The cortical slices were fixed in zinc-ethanol-formaldehyde (Korzhevskii *et al*., 2015), dehydrated, and embedded in paraffin. Paraffin blocks were cut into 5 μm sections. After the standard dewaxing procedure, the sections were unmasked by heating in modified citrate buffer S1700 (Dako, Glostrup, Denmark). Then, the activity of endogenous peroxidase was blocked in 3% hydrogen peroxide solution for 10 min. Then the tissue was blocked in Protein Block solution (Spring Bioscience, Pleasanton, USA) for 10 min. The sections were incubated for 48 hours in monoclonal (clone 3C12) mouse anti-Ezrin antibodies (1:100, Diagnostic BioSystems, Pleasanton, USA, catalog number Mob 380) in monoclonal (clone GA5) mouse anti-GFAP antibodies (RTU; Monosan, Netherlands, catalog number CM065). Reveal Compliment and Reveal HRP-conjugated solution from The Reveal Polyvalent HRP DAB Detection System kit (Spring Bioscience, USA) was used as a secondary reagent. The secondary antibody was applied for 30 min, according to the manufacturer’s instructions. Ezrin was visualized using 3,3’-diaminobenzidine from the DAB+ kit (Agilent, Santa Clara, USA). Then, all diaminobenzidine-developed samples for brightfield microscopy were mounted with the permanent medium Cytoseal 60 (Thermo Fisher Scientific). Samples were imaged with Leica DM750 microscope and an ICC50 camera (Leica, Germany). The mean immunofluorescence of Ezrin in the astrocyte territorial domain was normalized to the mean fluorescence of soma. The percentage of pixels with GFAP immunofluorescence exceeding a threshold was calculated in each frame.

### Western Blotting

Cortical tissue from the patients was frozen in liquid nitrogen. Then each tissue block was homogenized in the RIPA buffer containing SIGMAFAST protease inhibitor cocktail (Sigma-Aldrich, St. Louis, USA), diluted in the loading buffer (120 mM Tris-HCl, 20 % [v/v] glycerol, 10 % [v/v] mercaptoethanol, 4 % [w/v] sodium dodecyl sulfate, and 0.05 % [w/v] bromophenol blue, pH 6.8), loaded to SDS-PAGE, and blotted onto nitrocellulose membranes (GE Healthcare, Chicago, USA). The membranes were blocked in 5% skim milk (Sigma-Aldrich) in the TBS buffer (50 mM Tris, 150 mM NaCl, pH 7.4) + 0.2 % Tween-20 (Applichem, Darmstadt, Germany) for 1 h at room temperature and then incubated overnight at 4°C either with primary mouse antibodies for Ezrin (EZR, Antibodies-Online, catalog number ABIN5542456, Aachen, Germany), or with primary rabbit antibodies for GLT-1 (Abcam, Waltham, USA, catalog number 106289), or with primary rabbit antibodies for GFAP (Abcam, catalog number ABIN3044350), or with primary guinea pig antibodies for GS (Synaptic Systems, 367 005, Goettingen, Germany). After incubation with primary antibodies, membranes were rinsed in TBS with 0.2 % Tween-20 and incubated with corresponding HRP-conjugated secondary antibodies: either with anti-mouse IgG (Jackson Immunoresearch, West Grove, USA, catalog number 715-005-150), or anti-rabbit IgG (Abcam, catalog number 6721), or anti-guinea pig IgG (Jackson Immunoresearch, catalog number 706-035-048) for 1 h. ECL substrate (Bio-Rad, Hercules, USA) was used for signal detection. Protein bands were visualized using the ImageQuant LAS500 chemidocumenter (GE Healthcare, Chicago, USA). The ECL signal was visualized in the chemiluminescent channel, and the protein marker was detected in the optical channel of the chemidocumenter. The intensity of the protein bands was quantified using the gel analyzer option of ImageJ software (NIH, Bethesda, USA). To exclude inter-sample variability, the averaged intensities of the bands measured for each patient/protein were normalized to the average intensity of the lanes with the total protein of the corresponding samples using the No-Stain labeling kit (A44449, Life Technologies, Carlsbad, USA) using the fluorescent channel of the chemidocumenter.

### Statistics and reproducibility

Several cortical slices were collected from each patient. In patch-clamp recordings, only one cell was recorded per slice. N-number indicated the number of people in the sample, and n-number indicated the number of slices, cells, or samples used. Because of ethical considerations, no statistical method was used to determine the sample size. The sample size was determined by the availability of the access tissue. No assumption was made about the data distribution. Hence non-parametric Mann-Whitney statistical test was used. p > 0.05 considered non-significant (N.S.) The following values were considered significant *p < 0.05, **p < 0.01, ***p < 0.005. All data are presented as box and whiskers plots. Whiskers show 1.5 IQR. No data were excluded from the analyses, the experimenters were not blinded.

